# Genetic Basis of Melanin Pigmentation in Butterfly Wings

**DOI:** 10.1101/102632

**Authors:** Linlin Zhang, Arnaud Martin, Michael W. Perry, Karin R.L. van der Burg, Yuji Matsuoka, Antónia Monteiro, Robert D. Reed

**Author notes:** NCBI Gene Expression Omnibus (GEO): GSE78119. Corresponding authors: Department of Ecology and Evolutionary Biology, Cornell University, 215 Tower Rd., Ithaca, NY 14853-7202. (L.Z.); Department of Biological Sciences, National University of Singapore, 14 Science Drive 4, 117543, Singapore and Yale-NUS College, 6 College Avenue East, 138614, Singapore.sg (An.M.); Department of Ecology and Evolutionary Biology, Cornell University, 215 Tower Rd., Ithaca, NY 14853-7202. (R.D.R.).

## Abstract

Despite the variety, prominence, and adaptive significance of butterfly wing patterns surprisingly little known about the genetic basis of wing color diversity. Even though there is intense interest in wing pattern evolution and development, the technical challenge of genetically manipulating butterflies has slowed efforts to functionally characterize color pattern development genes. To identify candidate wing pigmentation genes we used RNA-seq to characterize transcription across multiple stages of butterfly wing development, and between different color pattern elements, in the painted lady butterfly *Vanessa cardui*. This allowed us to pinpoint genes specifically associated with red and black pigment patterns. To test the functions of a subset of genes associated with presumptive melanin pigmentation we used CRISPR/Cas9 genome editing in four different butterfly genera. *pale*, *Ddc*, and *yellow* knockouts displayed reduction of melanin pigmentation, consistent with previous findings in other insects. Interestingly, however, *yellow-d*, *ebony*, and *black* knockouts revealed that these genes have localized effects on tuning the color of red, brown, and ochre pattern elements. These results point to previously undescribed mechanisms for modulating the color of specific wing pattern elements in butterflies, and provide an expanded portrait of the insect melanin pathway.

## Introduction

Butterflies are a canvas upon which evolution paints with many varied hues. Indeed, for these highly visual creatures pigmentation is the primary medium used to communicate with other animals – whether to attract mates, to deter predators, or to escape detection. If we are to understand butterfly diversity we must work to better understand the origin and diversification of pigments themselves. Here we expand work on the genetic basis of butterfly pigmentation for several reasons. First, there is a large diversity of pigment types in butterflies, including both characterized and uncharacterized pigments, which provide an excellent model of how genetic biosynthetic mechanisms evolve in concert with physiology and morphology (NIJHOUT 1991). Second, several butterfly species are highly accessible laboratory models – they can be reared in large numbers, their wing tissues are large enough for functional genomics work, wings produce enough pigments for chemical characterization, and recently developed CRISPR/Cas9 genome editing methods now allow experimental validation of gene function (LI *et al*. 2015; PERRY *et al*. 2016; ZHANG AND REED 2016). Third, in many species pigments play important adaptive functions and show clear phylogenetic patterns of gain and loss, thus providing a link between developmental genetics and evolutionary biology. Finally, ongoing work by multiple research groups continues to define genes and regulatory networks involved in wing pattern evolution (BELDADE AND BRAKEFIELD 2002; KRONFORST AND PAPA 2015; MONTEIRO 2015; WALLBANK *et al*. 2016), thus providing a foundation for work aimed at understanding the upstream processes that regulate pigmentation genes. In this study we directly test the function of a suite of pigmentation genes in the painted lady butterfly, *Vanessa cardui*, the buckeye, *Junonia coenia*, the squinting bush brown, *Bicyclus anynana*, and the Asian swallowtail, *Papilio xuthus*. All of these are popular model species that display a wide range of many pigment and pattern types and are valuable systems for studying the evolution and development of wing patterns.

This study has three specific goals: (1) to characterize candidate genes implicated in ommochrome and melanin pigmentation using comparative transcriptomics, (2) to test if melanin genes characterized in other contexts in other insects (Figure 1) also play a role in butterfly wing pigmentation, and (3) to functionally characterize previously unknown genes potentially involved in pigmentation. To achieve these goals we employed a two-step process of transcript characterization followed by genome editing of candidate loci. Specifically, using RNA-seq in *V. cardui* we were able to identify both known and novel candidate wing pigmentation genes expressed during wing development. Using these sequences we then employed CRISPR/Cas9 genome editing to induce targeted deletions in eight selected candidate genes of the melanin synthesis pathway (MSP), which is associated with yellow, brown, and black pigment synthesis. Six of the eight targeted genes yielded wing pigmentation phenotypes across a total of four species. In addition to significantly improving our mechanistic understanding of butterfly wing pattern development, this work also expands our current knowledge of the insect melanin pathway. From a technical perspective, our work demonstrates how combining high-throughput transcript characterization with Cas9-mediated genome editing can be a powerful tool for discovering novel genes underlying morphological trait development in non-traditional model systems.

**Figure 1.**
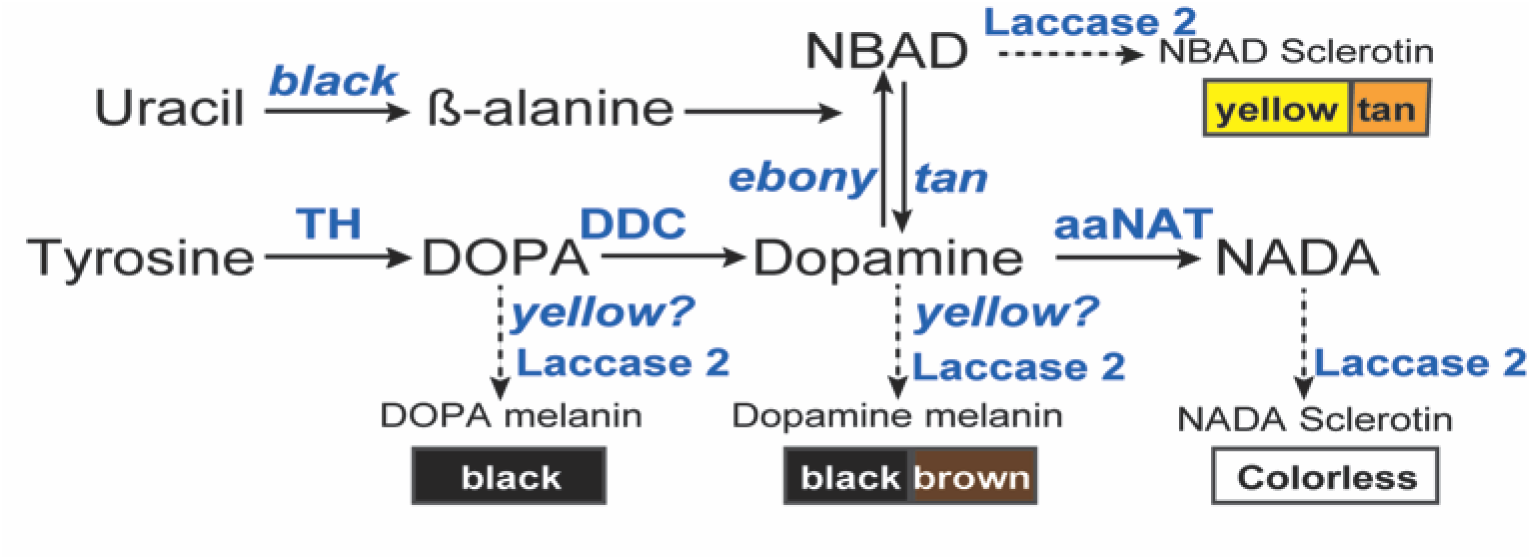
**Genetic model of the insect melanin biosynthesis pathway WRIGHT 1987)**. Tyrosine is the initial precursor for all insect melanins. Tyrosine hydroxylase (encoded by *pale*) and DDC (encoded by *Ddc*) convert tyrosine to dihydroxyphenylalanine (DOPA) and dopamine (dihydroxyphenylethylamine), which are the precursors for black melanin synthesis respectively. *yellow* is required for synthesizing DOPA and dopamine melanins, which are synthetic eumelanins. *Laccase 2* is a phenoloxidase gene required for cuticular pigmentation. Yellowish-tan hues are produced by N-β-alanyl dopamine (NBAD) sclerotin, which requires the function of *ebony*, *black*, and *tan*. N-acetyl dopamine (NADA), which is catalyzed by *Dat1*, is a major constituent of colorless or transparent cuticle.

## Materials and Methods

### Animals

*V. cardui* and *J. coenia* were maintained in a 16:8hrs light:dark cycle at 28 °C, *P. xuthus* were reared in a 14:10 light:dark cycle at 28°C, while *B. anynana* were reared at 27°C and 60% humidity in a 12:12 light:dark cycle. *V. cardui* butterflies originated from Carolina Biological Supplies, were fed on multi-species artificial diet (Southland Inc.), and induced to oviposit on leaves of *Malva parviflora*. *J. coenia* butterflies were derived from a laboratory colony maintained at Duke University (kind gift of Fred Nijhout), were fed on multi-species artificial diet (Southland Inc.) supplemented with dried leaves of the host plant *Plantago lanceolata*, and were induced to oviposit on leaves of *P. lanceolata*. *P. xuthus* were a line derived from wild-caught females from Kanagawa, Japan. Larvae were reared in the laboratory on fresh citrus leaves. *B. anynana*, originally collected in Malawi, have been reared in the lab since 1988. Larvae were fed on young corn plans and adults on mashed banana.

### RNA isolation and library construction

Hindwings from five different stages of *V. cardui* were sampled and used in this study including last instar larvae, 72h after pupation, 5d after pupation (pre-pigment pupae), ommochrome development (around 5.5d after pupation when red-orange ommochorome pigments started to be expressed) and melanin development (around 6d after pupation when black melanin pigments began to show up). Wings were rapidly dissected in PBS and stored in RNAlater at -80 °C (Life Technologies). Red and black regions from the melanin stage forewings were further separated as two samples (Figure 2A). RNA was extracted from each sample using Ambion Purelink RNA Mini Kit (Life Technologies). Due to the small size of the source tissue, each last instar larva wing disc sample consisted of hindwings taken from three individuals and pooled together in equal-molar amounts of RNA. For pupal wings, each sample represented RNA taken from a single individual. Two biological replicates were sampled for RNA-seq library construction from each developmental stage, resulting in 12 RNA-seq samples.

**Figure 2.**
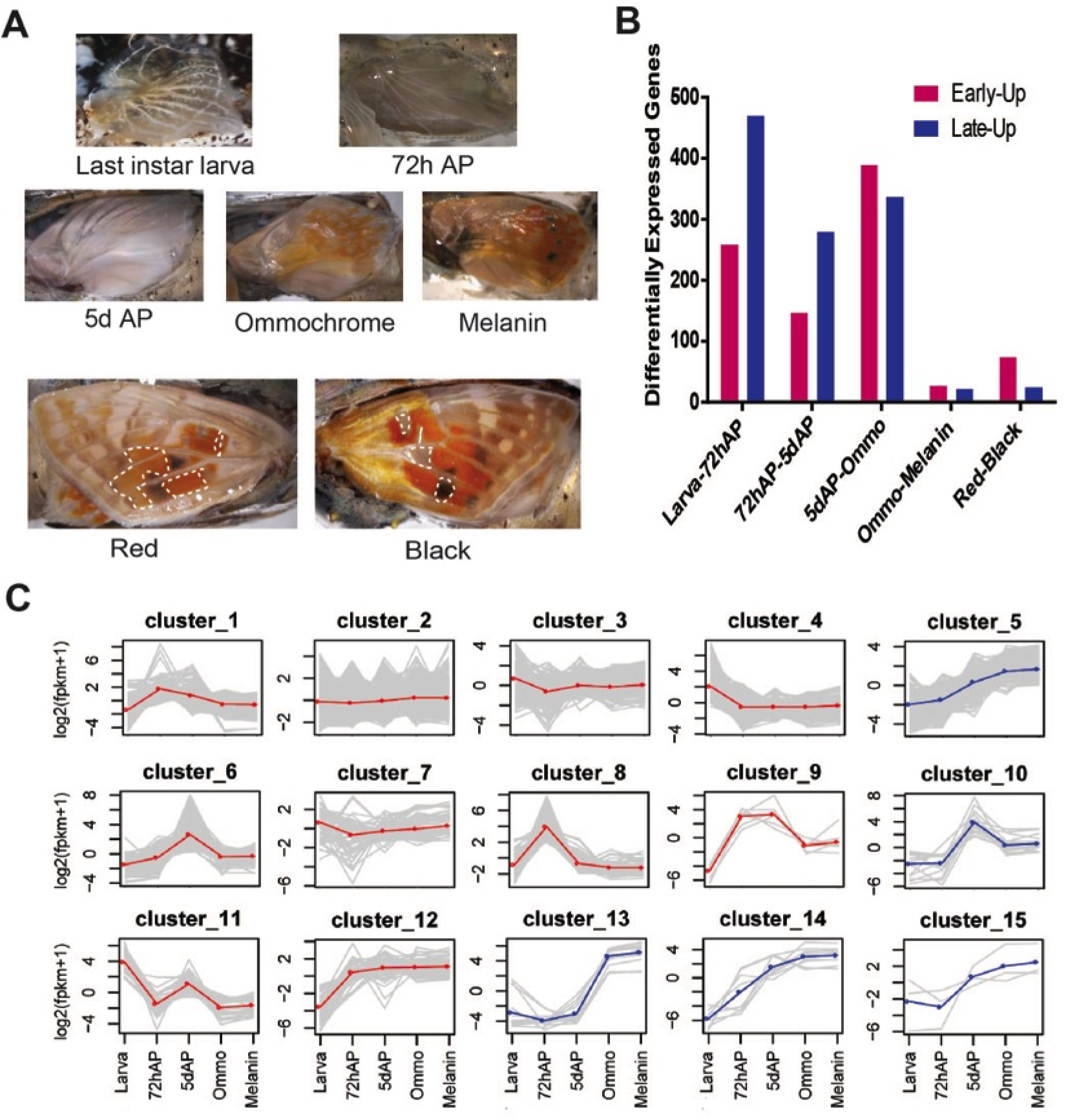
**RNA-seq analysis of the developing *V. cardui* butterfly wing transcriptome. (A)** The five developmental stages and two color regions sampled in this study (see Table S1 for details). “Black” and “Red” tissues were dissected at the melanin stage in forewings marked with white circles. **(B)** Number of differentially expressed unigenes in pairwise comparisons of consecutive stages (pink: upregulation in first, early stage); blue: upregulation in second, later stage. **(C)** Grouping of 13,875 unigenes into 15 clusters with similar expression profile. Clusters with upregulated expression patterns in later pupae are marked in blue.

Poly-A RNA was isolated with Oligo d(T)_25_ beads (New England Biolabs) from 1 μg total RNA of each sample and fragmented to size around 350-450 bp. First and second strand cDNA synthesis, end repair, 3′dA tailing and adaptor ligation were performed using NEBNext Ultra RNA library Prep kit for Illumina (New England Biolabs). All samples were tagged with a unique Illumina adapter index. The adapter-ligated sample was size-selected to ~500 bp fragments by AMPure XP beads (Beckman Coulter, Inc.). After 15 PCR cycles, libraries were quantified, multiplexed and sequenced on two Illumina Hiseq 2500 lanes with rapid run mode as 150 bp paired-end Illumina reads. Sequencing was performed at Cornell University’s Genomics Facility.

### Analysis of transcript expression data

After removing PCR primers, adapters, and low-quality reads, data sets from 12 samples were merged and an *in silico* normalization was performed to remove high coverage reads. Trinity (GRABHERR *et al*. 2011), which can take advantage of paired-end information, was used for assembly. The assembled sequences with putative isoforms identified by Trinity were filtered and only the longest isoform was kept. Cluster database at high identity with tolerance (CD-HIT) was used for further clustering and redundancy removal with minimum similarity cut-off of 95%. TransDecoder, a package included in the Trinity software, was used for prediction of coding regions. Assembled contigs were searched in the NR, Swissprot, and Flybase protein databases using BLASTX with a cutoff *E* value of 1e-5. Predicted proteins were further searched against InterProScan and Gene Orthology databases for domain and gene orthology annotation. TBLASTN and BLASTP packages were used to search for homologies with pigmentation genes in *D. melanogaster* as query sequences. Candidate genes were also aligned back to NR (protein) and NT (nucleotide) database for additional annotation.

Filtered reads from each sample were mapped to the assembly using Bowtie (LANGMEAD AND SALZBERG 2012). Gene expression levels for transcripts were measured by FPKM and normalized as suggested (CHEN *et al*. 2010). Differential expression analyses were performed for any pairwise comparison among five different developmental stages, last instar larva (Larva), 72 hours (72h AP), 5 days after pupation (5d AP), ommochrome (Ommo) and melanin developmental stages (melanin) using edge R with biological replicates and a cutoff FDR 0.001 (ROBINSON *et al*. 2010). Functional enrichment was further performed by identifying GO terms enriched in differentially expressed gene data sets (CHEN *et al*. 2010). We next conducted K-means clustering analyses to identify co-expressed genes during wing development. Clusters represented by genes highly upregulated in 5dAP, ommo, and melanin stages were extracted for further manual annotation. Pigmentation candidate genes were first identified by screening unregulated genes during pupal pigment maturation and then further categorized into two groups based on sequence homology and expression patterns. For example, ommochrome-associated genes are upregulated genes during pupal pigmentation stages that either orthologues of known ommochrome pathway genes or other ommochrome-rich color region specific expressed genes.

### Cas9-mediated genome editing

Our approach for producing G_0_ mosaic deletions in *V. cardui* and *J. coenia* butterfly wings followed the protocol of Zhang and Reed (ZHANG AND REED 2016). In general, we designed sgRNAs following the GGN_18_NGG or N_20_NGG rule on the sense or antisense strand of the DNA. Two sgRNAs, aiming to induce long deletions, were used for the target genes *Ddc*, *black*, *yellow*, *ebony*, *yellow-d*, *yellow-h2*, and *yellow-h3* in both *V. cardui* and *P. xuthus*, while one sgRNA, aiming to induce small indels, was used for *pale*, *yellow*, and *ebony* in *J. coenia* and *B. anynana* (Table S5). sgRNA template was produced by PCR amplification with a forward primer encoding a T7 polymerase binding site and a sgRNA target site, and a reverse primer encoding the remainder of the sgRNA sequence (BASSETT *et al*. 2013; BASSETT AND LIU 2014). sgRNAs were in vitro-transcribed using the MegaScript or MegaShortScript T7 Kits (Ambion) and purified by phenol:chloroform extraction followed by isopropanol precipitation or using MEGAClear columns (Ambion). Recombinant Cas9 protein was purchased from PNABio Inc. (catalog number CP01). We mixed 1μg of Cas9 protein and 375ng of each sgRNAs in a 5μl volume prior to injection. *V. cardui* eggs were collected from *Malva* leaves for 3 hours, lined up on double-sided adhesive tape onto a microscope slide and put into desiccant for 15 minutes. Slight variations in the method used across species can be found in Table S6. Microinjection of butterfly embryos was conducted using a pulled borosilicate glass needle (Sutter Instrument, I.D.:0.5mm) at an injection pressure of 20pps. Treated embryos were then incubated in a chamber at 28 °C and 70% humidity for the remainder of development. To confirm that our sgRNAs produce deletions at the desired target sites genomic DNA was extracted from a single caterpillar or legs of a butterfly adult by using proteinase K in digestion buffer (BASSETT *et al*. 2013). Fragments flanking the Cas9 target regions were amplified by PCR using primers flanking the deletion (Table S6). For the double sgRNA strategy, mutant PCR fragments were detected by agarose gel electrophoresis assay. For single sgRNA experiments, PCR fragments were detected with T7 endonuclease I (T7E1) assay as previously described (GUSCHIN *et al*. 2010).

In *B. anynana*, sgRNA target sequences were selected based on their GC content (around 60%), and number of mismatch sequences relative to other sequences in the genome (>3 sites). In addition we picked target sequences that started with a guanidine for subsequent in vitro transcription by T7 RNA polymerase. sgRNA templates were produced by PCR amplification [41, 42]. In vitro transcription of gRNAs were performed using T7 RNA polymerase (NEB) and purified by ethanol precipitation. We injected Cas9 protein (PNABio Inc.) or Cas9 mRNA as a source of CRISPR RNA-guided nuclease. We used the pT3TS-nCas9n vector (Addgene) for Cas9 mRNA in vitro synthesis. Template for Cas9 mRNA was generated by cutting the pT3TS-nCas9n vector with Xba. Capped Cas9 mRNA was in-vitro transcribed by using mMESSAGE mMACHINE T3 Kit (Thermo Fisher), and poly(A) tail was added by using Poly(A) Tailing Kit (Thermo Fisher). We co-injected 0.5 µg/µl final concentration of gRNA and 0.5 µg/µl final concentration of either Cas9 mRNA or protein into embryos within 1h after egg laying. Food dye was added to the injection solution for visualization. Injected embryos were incubated at 27 and 60% humidity. After hatching, larvae were moved to corn leaves, and reared at 27 with a 12:12h light:dark cycle and 60% relative humidity. To confirm that our sgRNAs produce deletions at the desired target sites genomic DNA was extracted from a pool of about 5 injected embryos that did not hatch with SDS and Proteinase K method, and used for T7 endonuclease (NEB) assay and sequence analyses. Primers for those analyses are listed on Table S6.

### Microspectrophotometry

Scale reflectance of wild-type and CRISPR mutants was measured at two specific regions (arrowheads in Figure 7A,C) of ventral forewings using an Ocean Optics USB 2000 spectrophotometer (Ocean Optics Inc.) attached to a compound light microscope.

### Imaging

Wing phenotypes were photographed using a Nikon DSLR camera equipped with a Micro-Nikkor 105mm – F2/8 Macro Lens, a VHX-5000 digital microscope, or a Zeiss Axiocam 506 color camera equipped with a Plan Apo S 1.0x FWD 60 mm Macro Lens on a SteREO Discovery.V20 microscope.

### Data Accession

***V. cardui*** wing RNA-seq raw reads on NCBI SRP062599. Full transcript assembly and expression profiles are available at NCBI Gene Expression Omnibus (GEO) database with accession number GSE78119. The transcript assembly and unigenes are also available for searching at *butterflygenome.org.*

## Results

### Transcriptome analyses

We used RNA-seq to generate a comprehensive profile of transcript abundance during wing developmental stages (last instar larva, 72h early stage pupa, 5d pre-pigment pupa, ommochrome and melanin development papa) and from different color patterns as a first step to investigate the molecular basis of wing pattern development in *V. cardui* (Figure 2A and Table S1). We first performed a *de novo* assembly based on a total of 174 million 150 bp paired-end Illumina Hiseq 2500 reads (GSE78119). The Trinity assembler generated 74,995 transcripts with longest ORF and a N50 length of 2,062 bp. 13,875 unigenes (“unigenes” are predicted transcripts, output from the assembly software), with a minimum length of 100 bp and a N50 length of 1,626 were predicted. A search against Swissprot and Flybase databases provided annotations for 13,064 and 13,636 unigenes based on an E-value of 1e-5. InterProScan and gene ontology (GO) analyses assigned 10,455 and 7,938 unigenes, respectively, into functional domains or categories.

We next aimed to identify general transcription patterns during wing development. Overall, 2,114 unigenes were found to be significantly differentially expressed (FDR<0.001) by pairwise comparison among the five stages (Figure 2B). Of these, 1,505 unigenes were upregulated by comparing consecutive stages, while 1,106 unigenes were downregulated. A total of 80 and 29 unigenes were identified to be specifically upregulated in red and black color regions, respectively. Final instar larvae (Larvae)-vs.-72h after pupation (72hAP) comparison showed the highest number of upregulated genes (n=673), perhaps reflecting major tissue state changes after metamorphosis. GO enrichment analyses of differential expression genes indicated dominant biological process during each representative developmental stage including larvae, early pupal, and pigmentation-stage pupal wings (Table S2). Interestingly, the day 5 stage, characterized by early development of wing scales, was enriched for the GO terms ‘structural constituent of cuticle’. We also note that the GO category ‘transporter activity’ was significantly enriched during the pigmentation phase. No functional term was enriched in comparison of ommochrome-stage and melanin-stage pupal wings (*P*>0.01).

K-means clustering of co-expressed genes revealed the dynamic nature of expression patterns throughout wing development (Figure 2C and Table S3). We divided expression series data into 15 clusters to identify genes with similar expression dynamics. Many sets of genes showed clear stage-specific expression patterns: clusters 3, 4, 7, and 11 showed the highest gene expression in last instar imaginal wing discs. Clusters 1 and 8 showed highest expression in early pupae, clusters 9, 12, and 14 showed highest expressed at 72h early pupae, and clusters 5, 10, 13, 14, and 15 showed highest expression in late pupal development, including at stages that show the visible deposition of pigments. These latter clusters include MSP genes such as *pale*, *Ddc*, *tan*, *black*, and *ebony* (Table S3), consistent with previous results (KOCH *et al*. 1998; FERGUSON *et al*. 2011b; HINES *et al*. 2012; DANIELS *et al*. 2014; CONNAHS *et al*. 2016). Of note, clusters 3 and 13, upregulated in both last instar larva and pigmentation stages, contained many known ommochrome pigmentation genes including *cinnabar* and *kynurenine formamidase*, consistent with previous results based on quantitative PCR in *V. cardui* (REED AND NAGY 2005). Overall, these clustering results show that blocks of genes involved in similar types of pigmentation tend to be regulated in similar temporal patterns.

### Identification of pigmentation candidate genes by RNA-seq

To identify genes potentially involved in wing pigment development, we screened transcript expression based on two criteria. First, we selected transcripts belonging to gene families previously implicated in pigmentation if they showed upregulation during pupal pigment maturation. Second, if a transcript was not related to a previously known pigmentation gene family, we selected it if it showed differential expression between red and black color regions, also during pupal pigment maturation.

*Ommochrome-associated genes:* We identified 26 genes associated with presumptive ommochrome pigment patterns, following the criteria described above (Table S4). All of the 26 genes were upregulated in late-stage red color patterns, and three were previously suspected to play a role in pigmentation. One of these, *optix*, is a transcription factor that promotes red wing patterns in *Heliconius* butterflies (REED *et al*. 2011; MARTIN *et al*. 2014). We also found high red-associated expression levels ofthe known ommochrome genes *cinnabar* and *kynurenine formamidase.* Many transcripts coding for poorly characterized transporters also showed red-associated expression patterns (Table S4). Importantly, seven major facilitator superfamily (MFS) transporter transcripts (LEMIEUX 2007) showed significant upregulation during pigment development. Of these, four showed strong and specific associated expression patterns with red regions. One transcript coding for an ATP-binding cassette (ABC) transporter C family member transcript also showed strong red-associated expression pattern (Table S4). Surprisingly, we also found two transcripts belonging to juvenile hormone binding protein family genes (HIRUMA *et al*. 1984) expressed in extremely strong association with ommochrome pigmentation. Another notable surprise was that the expression levels of the known *D. melanogaster* ommochrome transporters *scarlet* and *white* did not show differential expression patterns associated with pigmentation.

*Melanin-associated genes:* We identified 27 genes associated with presumptive melanin pigment patterns (Table S4), including six previously recognized melanin synthesis genes, all of the members of *yellow* gene family that have been previously characterized in other insects (FERGUSON *et al*. 2011a), and six transporters. Of these, 15 showed color pattern-specific expression where transcripts were upregulated in black vs. red patterns late in pupal development. We observed particularly strong upregulation of the melanin-associated genes *pale*, *Ddc*, *tan*, *ebony*, and *black* during both ommochrome- and melanin-stage pupae, with expression peaking at the onset of melanin pigment deposition (Figure 3A and Table S4). These results show that both ommochrome- and melanin-associated genes are activated at the ommochrome stage, however melanin gene expression persists longer. Similar to our results from the ommochrome candidates, we also observed three uncharacterized MFS transporter transcripts that showed significant upregulation in black patterns.

**Figure 3.**
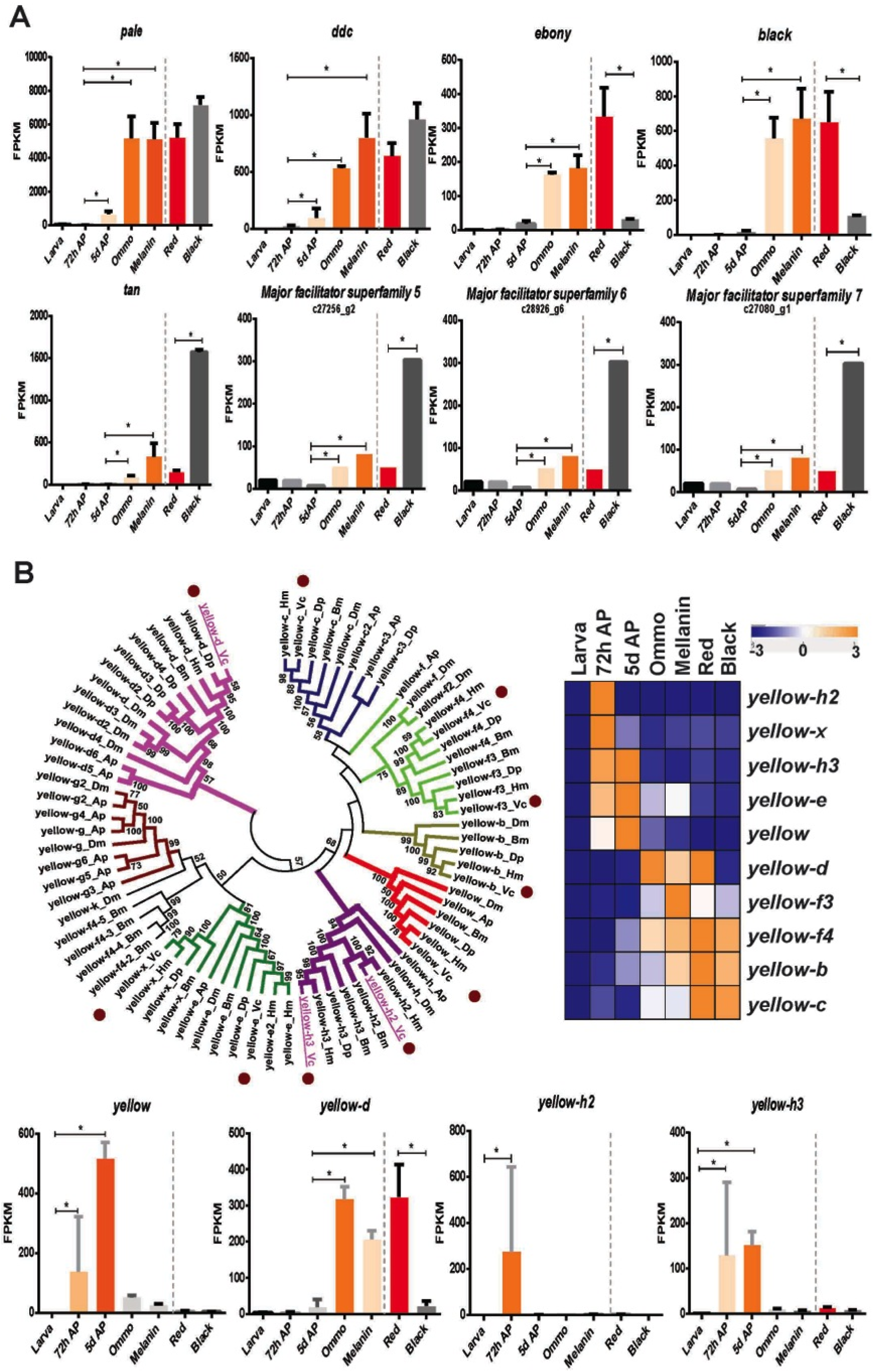
**Gene expression patterns of major genes of the melanin biosynthetic pathway in *V. cardui***. **(A)** Gene expression levels of known, major genes of the melanin biosynthetic pathway. The selected genes are those significantly differentially expressed during pupal pigment maturation and between black and red color patterns. (B) Phylogenetic analysis of insect *yellow* gene family members and their expression patterns in *V. cardui*. Neighbor joining tree of predicted yellow proteins from *Drosophila melanogaster* (Dm), *Acyrthosiphon pisum* (Ap), *Bombyx mori* (Bm), *Heliconius melpomene* (Hm), *Danaus plexippus* (Dp), and *V. cardui* (Vc). Putative orthologs in different species are denoted with the same color. Sequences from *V. cardui* are marked with brown circles. CRISPR targets are indicated in bold and underlined. The expression profiles of *V. cardui yellow* gene family members during different developmental stages as well as between red and black regions are indicated.

The *yellow* family of genes showed great diversity in expression profiles so we examined genes in greater depth. We identified ten *V. cardui yellow* family gene members by phylogenetic comparisons of protein sequences with those in *Heliconius melpomene*, *Danaus plexippus*, *B. mori*, *Acyrthosiphon pisum*, and *D. melanogaster* (Figure 3B). All ten genes showed upregulation during mid- or late-pupal development. Specifically, *yellow*, *yellow-e*, *yellow-h2*, *yellow-h3*, and *yellow-x* showed high expression level in pre-pigmentation early pupae, while *yellow-b*, *yellow-c*, *yellow-d*, *yellow-f4*, and *yellow-f3* showed higher expression during later pigment synthesis stages. Interestingly, *yellow-d* was observed with significantly higher expression level in red regions (*P*<0.001). These diverse expression patterns indicated that *yellow* gene family members might be involved in multiple previously uncharacterized aspects of butterfly pigment development.

### Functional validation of melanin pathway genes

We focused follow-up functional analyses on melanin biosynthetic pathway genes, in part because the functions of many of these genes are understood thanks to previous work in other insects (Figure 1), and thus provide clear predictions for potential phenotypes. We selected a total of eight candidate melanin genes for functional assessment by CRISPR/Cas9: *pale*, *Ddc*, *ebony*, *black*, *yellow*, *yellow-d*, *yellow-h2*, and *yellow-h3* (Table S5). This represents a range of genes that would be predicted to have a general negative effect on melanization (*pale*, *Ddc*, and *yellow*), which should alter melanin level and/or type (*ebony* and *black*), or are of unknown function (*yellow-d*, *yellow-h2*, and *yellow-h3*). As described below, six out of eight gene loss-of-function experiments yielded clear G_0_ generation mosaic phenotypes, and PCR-amplified fragment length genotyping validated the efficacy of our approach for producing deletion mutations at the targeted sites (Table S5-6 and Figure S1).

***pale***: *pale* encodes the tyrosine hydroxylase (TH) enzyme which catalyzes the formation of DOPA from tyrosine at the earliest metabolic step of the melanin synthesis cascade (Figure 1). We produced *pale* deletions in *J. coenia* using a single sgRNA. *pale* deletions resulted in severe cuticular defects including improperly expanded wings, and patches showing failed scale development on the thorax and wings (Figure 4A-E). Tyrosine hydroxylase deficiency also triggered complete amelanism in scales that emerged properly, with effects varying depending on their ground state: in the thorax, mutant scales shifted from dark brown to white (Figure 4B); most wing mutant scales showed a lighter coloration and an altered morphology such as a rounded aspect, instead of a marked scalloping (Figure 4F-F’); and finally, melanic scales of the eyespot inner rings became completely transparent, also taking a disheveled aspect due to an apparent structural defect or thickness reduction (Figure 4F-F’). Interestingly, mutant eyespots also lost the characteristic blue color of their foci (central dots, Figure 4G-H), and inspection at high-magnification reveals that this is associated with a transformation of black ground scales into transparent scales. In other words, the loss of the blue hue is not due to a structural defect in blue scales; rather, the cover scales responsible for the blue coloration lack a black background in the mutant context that is necessary for blue reflectance by super-imposition, an effect described in blue-iridescent *Morpho* butterflies (GIRALDO AND STAVENGA 2016).

**Figure 4.**
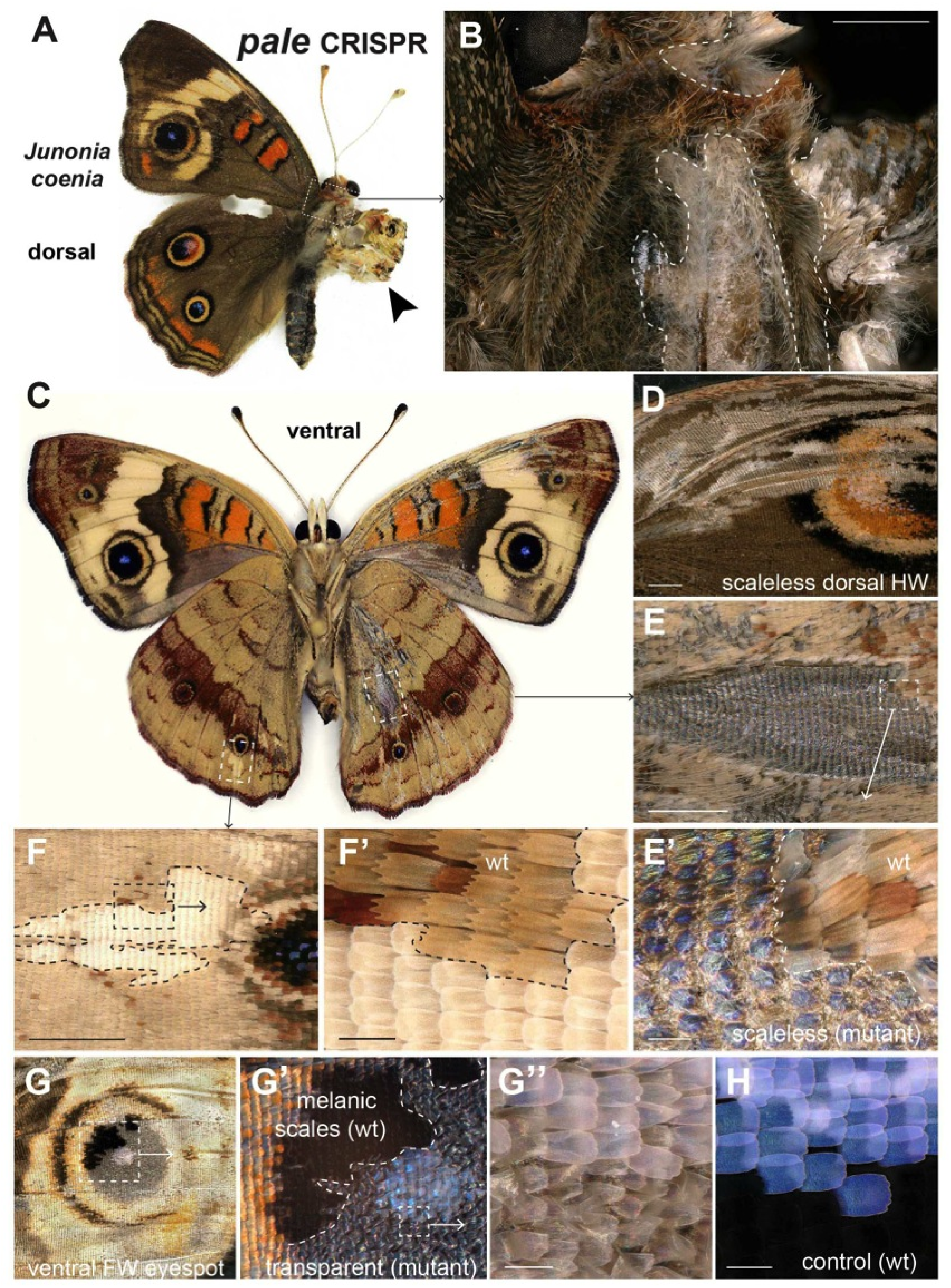
**Effects of *pale* somatic mutagenesis in *J. coenia.* (A-B)** Mosaic *pale* mutant showing severe one-sided defects in sclerotization and pigmentation (dorsal view), including improper wing expansion (arrowhead), and missing or amelanic scales (dotted lines). **(C-F’)** Somatic *pale* mutant showing varied effects on scale morphology (**C**, ventral view; **D**, dorsal hindwing), including scaleless patches **(D–E)**, and depigmented round scales lacking scalloped ridges **(F-F’)**. **(G-G”)** Mutant ventral forewing eyespot showing the transformation of thin transparent scales. Residual black color in the mutant patch is due to the opposite wing surface. The white color of the mutant eyespot focus compared to the wild-type blue focus **(H)** is due to the transformation of black ground scales into transparent scales. Scale bars: 1000μm (**B**, **D-F**); 100μm (**F’**, **E’**, **G”**, **H**).

***yellow***: *yellow* encodes a secreted extracellular protein required for production of black melanin pigments in *Drosophila* (BIESSMANN 1985) (Figure 1). We produced *yellow* deletions in *V. cardui*, *B. anynana*, and *P. xuthus* using single or double sgRNAs. *yellow* deletion mosaics revealed strong reduction of presumptive melanins in both *V. cardui* and *B. anynana* wings, and in *P. xuthus* wings as previously reported (PERRY *et al*. 2016) (Figure 5A). In *B. anynana* overall brown regions became yellow, and the black scales became brown. We also observed marked larval phenotypes in *V. cardui* and *B. anynana*, including larvae with depigmented head capsules (Figure 5B). Unlike for *pale, yellow* deletions appeared to have no obvious effect on red wing scales, thus suggesting a primary role for this gene in the synthesis of black melanin pigments, with little or no effect on scale cuticularization or non-melanin pigment deposition.

**Figure 5.**
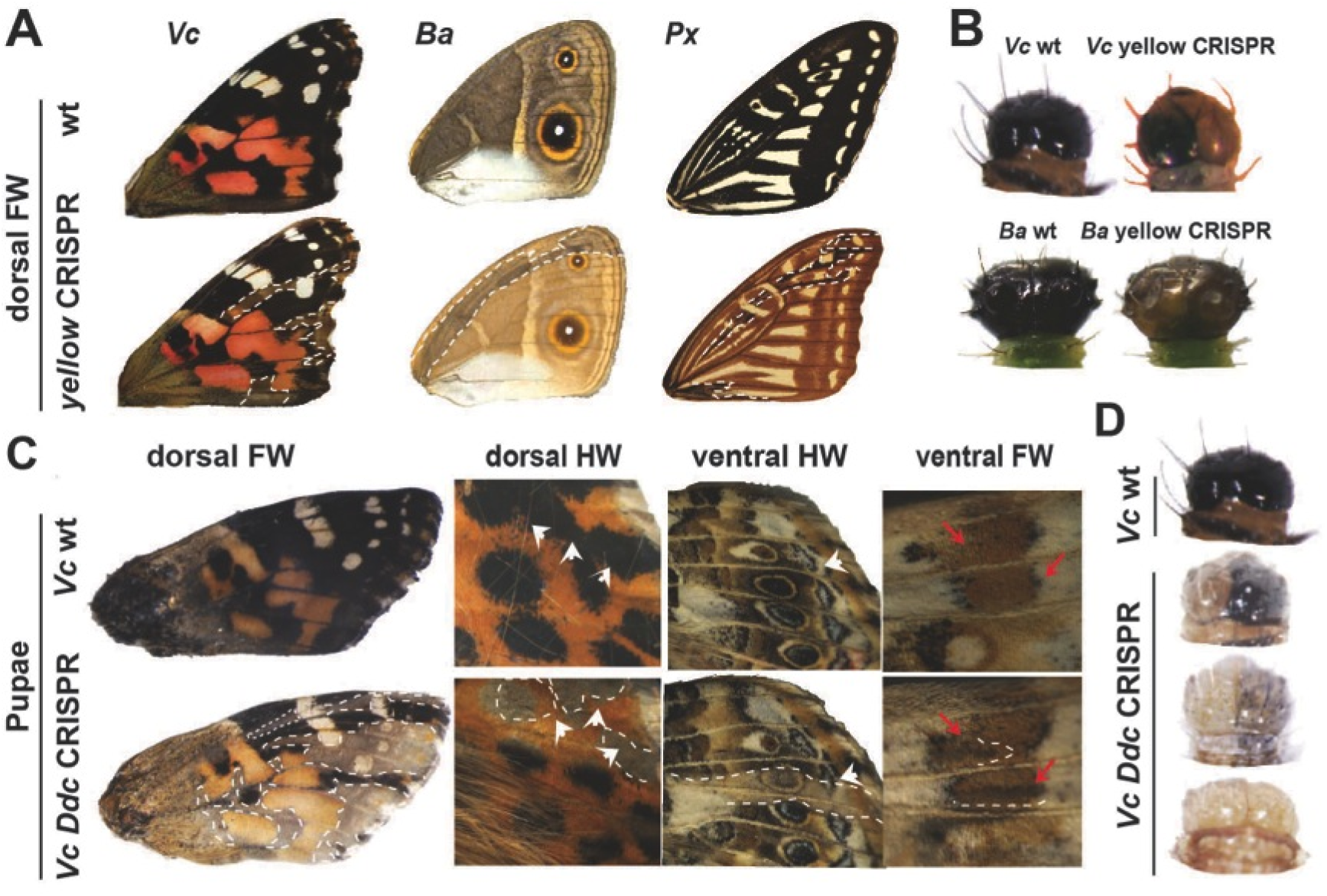
***yellow* and *Ddc* loss-of-function induces wing depigmentation mutants.** (A) Dorsal view of forewings in wild type and *yellow* deletion butterflies (left-right: *V. cardui*, *B. anynana*, and *P. xuthus*). Dotted lines: mosaic depigmentation phenotypes. (B) *yellow* deletion result in depigmentation of larvae head in *V. cardui* and *B. anynana*. (C) Examples of *Ddc* deletion *V. cardui* phenotypes in black region of dorsal forewing, black eyespots and parafocal elements of dorsal hindwing (white arrowhead), colored rings of ventral hindwing eyespot (white arrowhead), and grey strip of ventral forewing (red arrow). (D) Different levels of depigmentation in mosaic larval heads of *V. cardui*.

***Ddc***: *Ddc* encodes the melanin synthesis enzyme dopa-decarboxylase (Figure 1). We produced *Ddc* deletions in *V. cardui* using double sgRNAs. As previously reported (ZHANG AND REED 2016), *Ddc* deletions produced a phenotype similar to *yellow* with strong depigmentation of black scales and no effect on red scales (Figure 5C, D). Unlike for *yellow*, however, *Ddc* deletions also changed brown and tan pigments on the ventral surface (Figure 5C). *Ddc* knockout larvae were typically unable to break through their eggshells, perhaps due to an incomplete sclerotization of their mouthparts as seen in *D. melanogaster* mutants (WRIGHT *et al*. 1976b). For this particular locus it is important to note the caveat that knockouts resulted in such high mortality rates that we generated only two adults with wing phenotypes. We did, however, observe a large number of larvae with reduced melanin phenotypes (Figure 5D), providing clear evidence of *Ddc*’s role in melanin synthesis.

***ebony***: *ebony* encodes the N-β-alanyldopamine (NBAD) synthase enzyme which promotes the formation of light-color NBAD pigment derivatives (Figure 1). We produced *ebony* deletions in *V. cardui* using double sgRNAs, and *J. coenia* and *B. anynana* with a single sgRNA. Generally, *ebony* deletions lead to strong darkening of wing phenotypes, with some interesting stage-dependent nuances (Figure 6A-D). Late-stage pupal wings from *V. cardui* with *ebony* deletions showed marked, dramatic phenotypes where red (e.g. central band), white (e.g. eyespots), and brown (e.g. marginal band) color patterns turned black, producing an almost inverse coloration of the wild type wing (Figure 6A-D). We did not recover this phenotype in fully-emerged adult *V. cardui*, leading us to speculate these very strong phenotypes resulted in pupal mortality before normal melanin pigment development. In fact, *ebony* deletions resulted in many pupae being hyperpigmented (Figure 6E). This excessive melanization of internal tissues reflects *ebony* deficiency across large somatic clones, which would be predicted to have multiple deleterious effects on nervous system function, based on *Drosophila* knockout phenotypes (PÉREZ *et al*. 2010). We did, however, recover adults with darker mutant clones in both *B. anynana* and *V. cardui* (Figure 6B, F). In *B. anynana*, the brown background color (asterisk) was unaffected, but the white band regions, and the yellow rings of the eyespots became brown (Figure 6F). We also recovered mosaic *J. coenia* adults that showed *ebony* mutant clones consisting of dark scale patches (Figure 6G-G’’). The strength was variable across a wing surface, probably reflecting different allelic dosages between clones (Figure 6G’). In all *J. coenia* (Figure 6H-I), *V. cardui* (Figure 6D), and *B. anynana* (Figure 6F) these hypermelanic states affected red-orange (e.g. Basalis, Discalis I and II elements in *J. coenia* and central band in *V. cardui*) and buff/yellow scales (e.g. dorsal hindwing eyespot ring in *J. coenia* and ventral band and eyespot rings in *B. anynana*) that are traditionally considered to reflect an ommochrome composition (NIJHOUT AND KOCH 1991). Interestingly, we found no effect of *ebony* knockouts on the *J. coenia* dorsal brown background color field (asterisk, Figure 6I’) and *B. anynana* ventral brown background color field scales (asterisks, Figure 6F) in spite of predictable mutant clone extensions into these wing territories (Figure 6I-I’, and Figure 6F). It is thus possible that this gene is transcriptionally inactive in the corresponding scale cell precursors. In contrast, because mutant scales of the yellow eyespot rings of the dorsal surface in *J. coenia* show a similar color to the brown background (Figure 6I”), we speculate that *ebony* expression and NBAD synthesis are important for making this inner ring lighter than the dark color on the rest of this wing surface.

**Figure 6.**
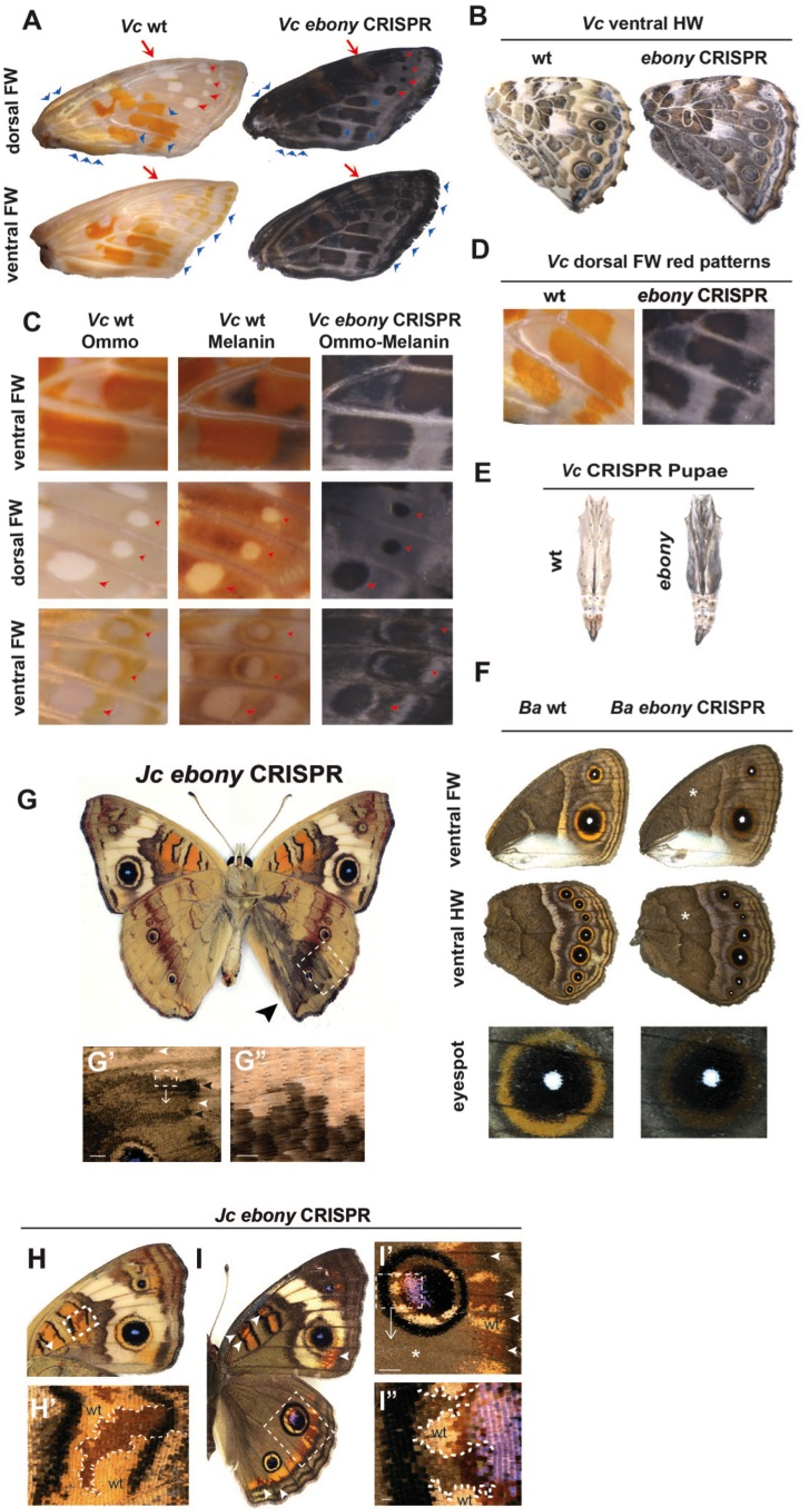
***ebony* loss-of-function results in hyperpigmented wing phenotypes. (A)** Dorsal and ventral view of forewings in wild type and *ebony* deletion pupal wings in *V. cardui*. Obvious patches of mutant tissues can be observed (arrow and arrowhead). (B) Ventral view of hindwings in wild type and *ebony* deletion adults in *V. cardui*. **(C)** Detailed morphology of forewings reveals wing scales turn black at the initiation of ommochrome stage in *ebony* deletion *V. cardui* mutant. Arrowhead: obvious mutant tissues. (**D**) Red colors are disturbed in *V. cardui ebony* deletion mutants. (**E**) *V. cardui* wild type and *ebony* deletion pupae. (**F**) Ventral view of wild type and *ebony* deletion adults in *B. anynana*. Darker scales were observed in the yellow ring of *B. anynana* eyespots in *ebony* deletion mutants. **(G-G”)** Mosaic hypermelanization of a ventral hindwing following *ebony* somatic knockout in *J. coenia.* Variable darkness of the mutant clones may reflect different allelic states (black and white arrowheads in **G”**) Scale bars: 500μm (**F’**) and 100μm (**F”**). **(H-H’)** Ventral forewing mutant clones showing darkening of orange patterns in *J. coenia* (arrowheads and dotted lines). **(I-I”)** Effects of *ebony* deletion on dorsal patterns in *J. coenia*, including the darkening of orange patterns (arrowheads), and of the eyespot inner ring (dotted lines). Mutant scales of the yellow ring resemble the brown background scales (asterisk), which are not affected by *ebony* deletion. Scale bars: 100μm (**I”**), 1000μm (**H’**, **I’**).

***black***: *black* encodes a protein for synthesizing β-alanine, which is enzymatically conjugated to dopamine to form NBAD (Figure 1). We produced *black* deletions using double sgRNAs in *V. cardui.* We found that *black* deletions caused darkening of a subset of color fields on the ventral wings (Figure 7A, B), especially the distal ventral forewing (Figure 7C). The hypermelanic state affects most of the brown and buff-colored pattern elements including strip between stripe between MI and EIII, marginal elements, forewing wing eyespot yellow rings (Figure 7A1-A2), hindwing basal, and hindwing central regions. To investigate the quantitative nature of the pigmentation difference, we measured reflectance spectra from ten individual scales from the same regions of both wild type and *black* knockout butterflies (Figure 7C). We found significant changes of reflectance spectra, with peak reflectance at 575-625 nm in wild type and 600-650 nm in *black* mutants. *black* mutants also showed a dramatic decrease in light reflectance, consistent with an overall darkening of color. We also recovered hyperpigmented pupae (Figure 7D).

**Figure 7.**
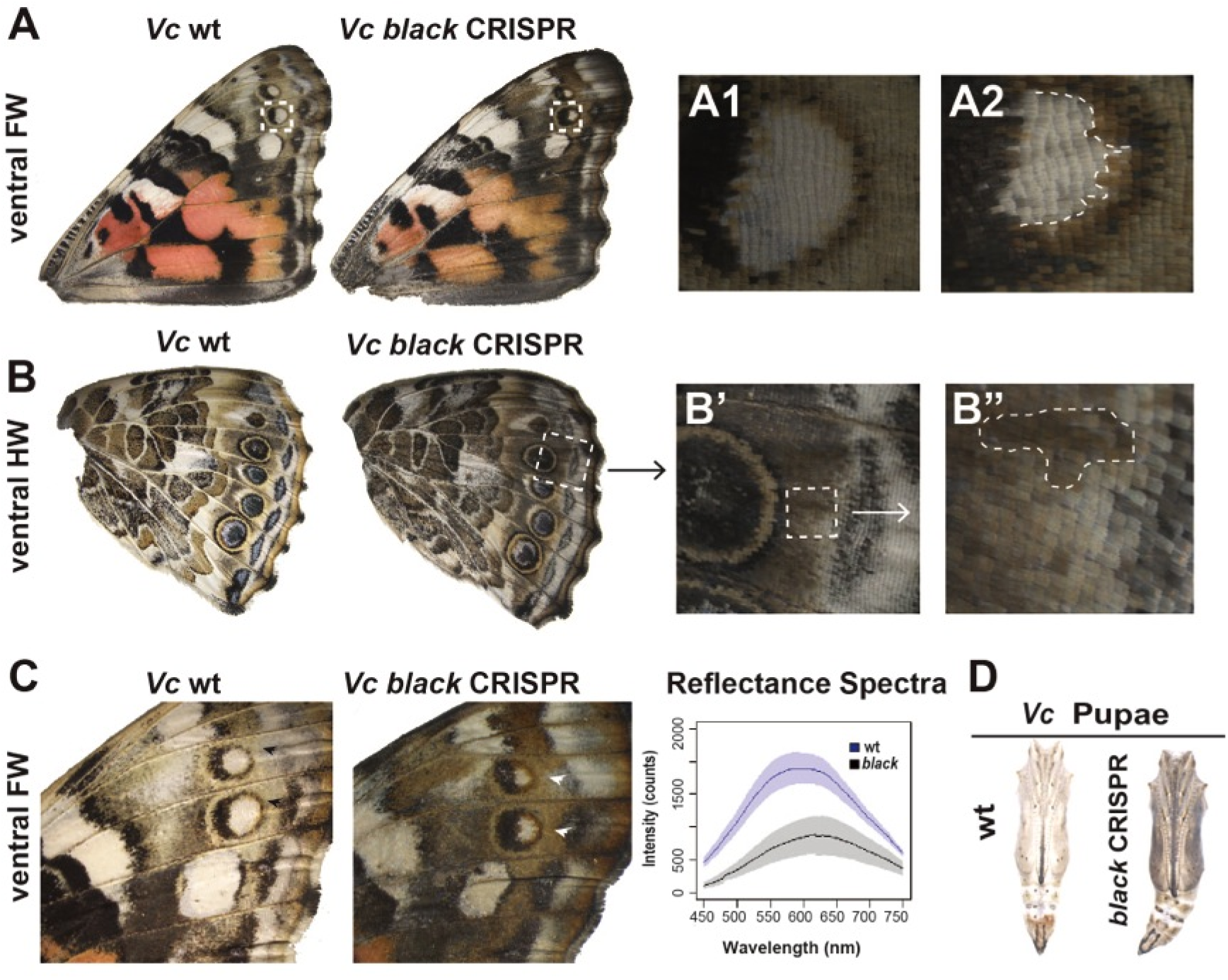
**Wing pattern-specific effects of *black* loss-of-function in *V. cardui*.** (**A**) Ventral view of wild type and *black* deletion wings in *V. cardui* shows enhanced melanization phenotypes. Darker yellow ring of eyespot (**A2**) can be observed compared to wild type (**A1**). (**B-B”**) *black* deletion causes hypermelanization of a ventral hindwing in *V. cardui.* (**C**) *black* deletion in *V. cardui* results in ectopic black pigments. Graphs of reflectance spectra of ventral forewing (white arrowhead) show significant shift in coloration between wild type and CRISPR mutants. (**D**) *black* deletion results in hyperpigmented pupae in *V. cardui*.

***yellow-d***: *yellow-d* encodes a protein of unknown function. We produced *yellow-d* deletions using double sgRNAs in *V. cardui.* We observed that *yellow-d* deletion promoted orange-brown pigmentation in specific ventral forewing patterns, especially the submarginal and marginal elements between EI and EII, and the stripe between MI and EIII including the rings around border ocelli (Oc) (Figure 7A). Optical reflectance spectra of affected pattern elements on the ventral forewing show significant differences in absorbance between wild type butterflies and *yellow-d* knockouts, as well as subtle difference in shape of spectrum (Figure 7B). Mosaic orange pigmentation was also found in ventral hindwings (Figure 7C-C’’), e.g. marginal elements between EI and EII, eyespot rings, and Discalis I element), and red patterns in dorsal forewings (Figure 7D-D’’).

***yellow-h2*** and ***yellow-h3***: *yellow-h2* and *yellow-h3* encode proteins whose transcripts have shown interesting color-related expression patterns in other species, however are of poorly understood function (FERGUSON *et al*. 2011a; FUTAHASHI *et al*. 2012). We produced *yellow-h2* and *yellow-h3* deletions using double sgRNAs in *V. cardui.* All injected animals that survived to pupation (15 *yellow-h2* and 26 *yellow-h3* pupae) died during late pupal pigment development. PCR genotyping validated the existence of long deletions in all dead pupae (Figure S1), however detection of possible pigment phenotypes was confounded by tissue necrosis.

## Discussion

In this study we significantly expand our knowledge of butterfly wing pattern development by functionally characterizing six genes underlying wing pigmentation. Our strategy was to use comparative RNA-seq to identify candidate pigment gene transcripts, and then to use Cas9-mediate targeted deletions to test the function of a subset of these candidates. RNA-seq is widely employed in non-model organisms to characterize candidate genes involved in the development of interesting traits, however such studies are often frustrated by long lists of candidate genes that are difficult to interpret due to a lack of functional validation. Our work shows how combining comparative transcriptomics and genome editing can provide a powerfully synergistic approach for exploring the evolution and development of traits in emerging model systems.

Our comparative transcriptomic work identified over 2,000 transcripts showing significant expression differences between wing development stages and color patterns. Using stringent screening criteria, we identified 26 candidate ommochrome-associated genes and 27 candidate melanin-associated genes, complementing previous wing transcriptome studies (FUTAHASHI *et al*. 2012; HINES *et al*. 2012; DANIELS *et al*. 2014; CONNAHS *et al*. 2016) by providing a more detailed portrait of pigment-specific expression patterns in *V. cardui*, a species that is highly amenable to genome editing. Although we will focus most of our discussion below on functional results, there are two surprising expression results we would like to highlight. First, finding that *optix* is upregulated in red patterns in *V. cardui* contradicts conclusions from previous comparative work at earlier stages of pupal development that suggested that the red color pattern-related expression of this gene was limited to heliconiine butterflies (REED *et al*. 2011; MARTIN *et al*. 2014). Our new results imply that *optix* may indeed play a color patterning role at later stages of development in non-heliconiine butterflies, although confirmation of this will require further work. Second, we identified a number of MFS transporters that showed highly specific associations with either red or black color patterns, suggesting that these genes might play roles in both ommochrome and melanin pigmentation. Indeed, MFS genes have been previously associated with insect ommochrome biosynthesis in *Bombyx* eggs, eyes, and larvae (OSANAI-FUTAHASHI *et al*. 2012; ZHAO *et al*. 2012), and melanin pigmentation in *Bombyx* larval cuticle (ITO *et al*. 2012). Interestingly, however, these previously characterized MFS genes are not orthologous to the ones identified in this study. Therefore, we speculate that many members of the diverse MFS gene family may play roles in the development of different pigment types across insects. We are currently working to better characterize these, and related genes.

Taking advantage of our transcriptome data we were able to use Cas9-mediated genome editing to test the wing pigmentation roles of eight candidate melanin synthesis genes. Of these, *pale*, *yellow*, and *Ddc* are previously known as melanin-promoting factors. CRISPR deletions in these three genes produced melanin repression phenotypes in accordance with expectations from *Drosophila* (MORGAN 1916; WRIGHT *et al*. 1976a; TRUE *et al*. 1999; WITTKOPP *et al*. 2002b) and other insects including *Papilio xuthus* (FUTAHASHI AND FUJIWARA 2005), *Manduca sexta* (GORMAN *et al*. 2007), *Bombyx mori* (FUTAHASHI *et al*. 2008; LIU *et al*. 2010), *Tribolium castaneum* (GORMAN AND ARAKANE 2010), and *Oncopeltus fasciatus* (LIU *et al*. 2016). Our findings support the idea that *pale*, *yellow*, and *Ddc* play deeply conserved roles insect melanin pigmentation. The structural defects we observed in *pale* knockout butterflies are also consistent with the known role of DOPA in sclerotization (ANDERSEN 2010). It is worth noting that *yellow* mutants in *V. cardui*, *P. xuthus*, and *B. anynana* all resulted in widespread defect clones in both forewing and hindwing, suggesting potential similar involvement of *yellow* between forewing and hindwing. This is consistent with findings in *Drosophila* but in contrast with *Tribolium* and *Oncopeltus*. In *Oncopeltus* RNAi knockdown of *yellow* caused much greater defects in hindwing than forewing (LIU *et al*. 2016), and only hindwing defects (not elytra) were observed in *Tribolium* RNAi knockdowns (ARAKANE *et al*. 2010). Such differences indicate that region-specific deployment of *yellow* might have changed through insect evolution. In sum, our findings support a growing body of literature to suggest that there is a highly conserved set of genes involved in insect melanin pigmentation, and that these genes have been deployed in multiple tissue types throughout evolutionary history.

Our results with *yellow-d*, *ebony*, and *black* were of particular interest to us because they showed that these genes can affect the hue of color patterns that do not appear to be typical black dopa-derived eumelanins. One of the most novel findings from our study relates to the function of *yellow-d*, which is a protein of previously uncharacterized function. Previous work has shown an association between *yellow-d* expression and dark pupal cuticle in *B. mori* (XIA *et al*. 2006), and red wing patterns in *Heliconius spp.* (FERGUSON *et al*. 2011a; HINES *et al*. 2012), but thus far there had been no functional evidence for a role in pigmentation, and these previous associations were potentially contradictory. We were thus interested to find that in *V. cardui* deletion of *yellow-d* resulted in specific buff-colored color pattern elements converting to a chocolate-ochre hue (Figure 7C, 8B). Although determining the chemical basis of this color variation is beyond the scope of this study, it is notable that this hue is not one that is normally associated with eumelanin pigments. We tentatively speculate that this color may be generated by a pigment or pigments in the NBAD sclerotin or pheomelanin pathways, and/or even perhaps ommochromes, although additional work will be required to confirm this. Unlike the other yellow gene family member *yellow*, loss of *yellow-d* function not only affect melanin patterns but also presumptive ommochrome patterns. This finding is consistent with the different expression profiles of these genes, where *yellow-d* shows red-specific expression while no differential expression was detected for *yellow*. We further speculate that co-expression of *yellow-d* and the other two melanin-suppressing genes *ebony* and *black* (Table S4) may indicate that these genes can be recruited together in to color-tune specific pattern elements. In any case, our *yellow-d* deletion results clearly demonstrate that the activity of this single gene is sufficient to tune the color of specific pattern elements by switching pigments.

Our *ebony* results were also of significant interest, again due to the dramatic effect of *ebony* deletions on non-eumelanin wing patterns. Work in *Drosophila* has shown that *ebony* suppresses melanin development in some tissue types, presumably because the enzyme encoded by *ebony,* NBAD-synthase, depletes dopamine that would otherwise be used for eumelanin synthesis. Therefore, *Drosophila ebony* mutants show an overall increase in melanization, while ectopic expression of *ebony* inhibits melanization (WITTKOPP *et al*. 2002a). Supporting this, *ebony* mutagenesis in *P. xuthus* showed enhanced melanic pigmentation in larvae (LI *et al*. 2015), and dark wing patches in *B. anynana* (BELDADE AND PERALTA 2017). Based on these previous results we were therefore unsurprised to see darkened wing clones in adults of *V. cardui*, *B. anynana*, and *J. coenia* (Figure 6). Interestingly, however, this darkening occurred across a range of color pattern types in all species – including buffs, tans, and light and dark browns. The most marked deletion phenotype was seen in a subset of *V. cardui* pupae we dissected, where developing pupal wings showed an almost complete conversion of all non-melanic color patterns to black, including red ommochrome and colorless (white) pattern. These results strongly suggest that *ebony* is a strong repressor of melanin synthesis in all wing scales, regardless of specific pigment type. *ebony* deletion in ommochrome stage wings caused darkening phenotypes in the scales that would become black in the melanin stage (Figure 6A), and we know that *ebony* is significantly upregulated at the ommochrome stage. Together these findings suggest that *ebony* suppresses melanin synthesis in black patterns before black pigmentation is produced. Furthermore, the phenotype of increased melanin in red scales correlates with the transcription profile where *ebony* showed red-specific expression. This is consistent with previous speculation based on correlations between *ebony* expression and red ommochrome color patterns in *Heliconius spp.* (FERGUSON *et al*. 2011b; HINES *et al*. 2012). These results are also consistent with *Oncopeltus ebony* RNAi knock down phenotypes in forewings, where *ebony* appears to suppress melanization in the orange pattern. The aberrant red patterns in *ebony* CRISPR knockout butterflies are known to show strong incorporation of ommochrome-precursor tryptophan (NIJHOUT AND KOCH 1991). We notice, however, that radiolabeling experiments in *J. coenia* also show weak levels of tyrosine incorporation in these patterns. Our current hypothesis is thus that orange scales synthesize light-color melanin derivatives such as NBAD sclerotin, and that *ebony* loss-of-function triggers the accumulation of dopamine, which is in turn converted into dark melanin (Figure 1). It is also plausible that an unidentified type of tyrosine-tryptophan hybrid pigment, such as a papiliochrome, may occur in *J. coenia*, although further work would be required to test this.

A third gene that had notable effects on wing coloration was *black*. Disruption of this gene is known to result in β-alanine deficiencies, therefore limiting NBAD sclerotin production to result in an excess of dopamine. This excess dopamine is thought to underlie the mutant phenotypes observed in various insects – an overall increase in cuticular melanization (HORI *et al*. 1984; ROSELAND *et al*. 1987; WRIGHT 1987). In butterflies we also observed similar enhanced melanization phenotypes in wing scales and pupal cuticle. While the apparently conserved function of *black* may not be remarkable by itself, the wing pattern specificity of the effect is indeed notable. Specifically, *black* deletions seemed to have a highly specific effect on the same color pattern elements that *yellow-d* affected, and resulted in a similar darkening of hue, though without the marked yellow overtones (Figure 8). These results tentatively suggest that local activity of *black* may be implicated in color-tuning specific color pattern elements.

**Figure 8.**
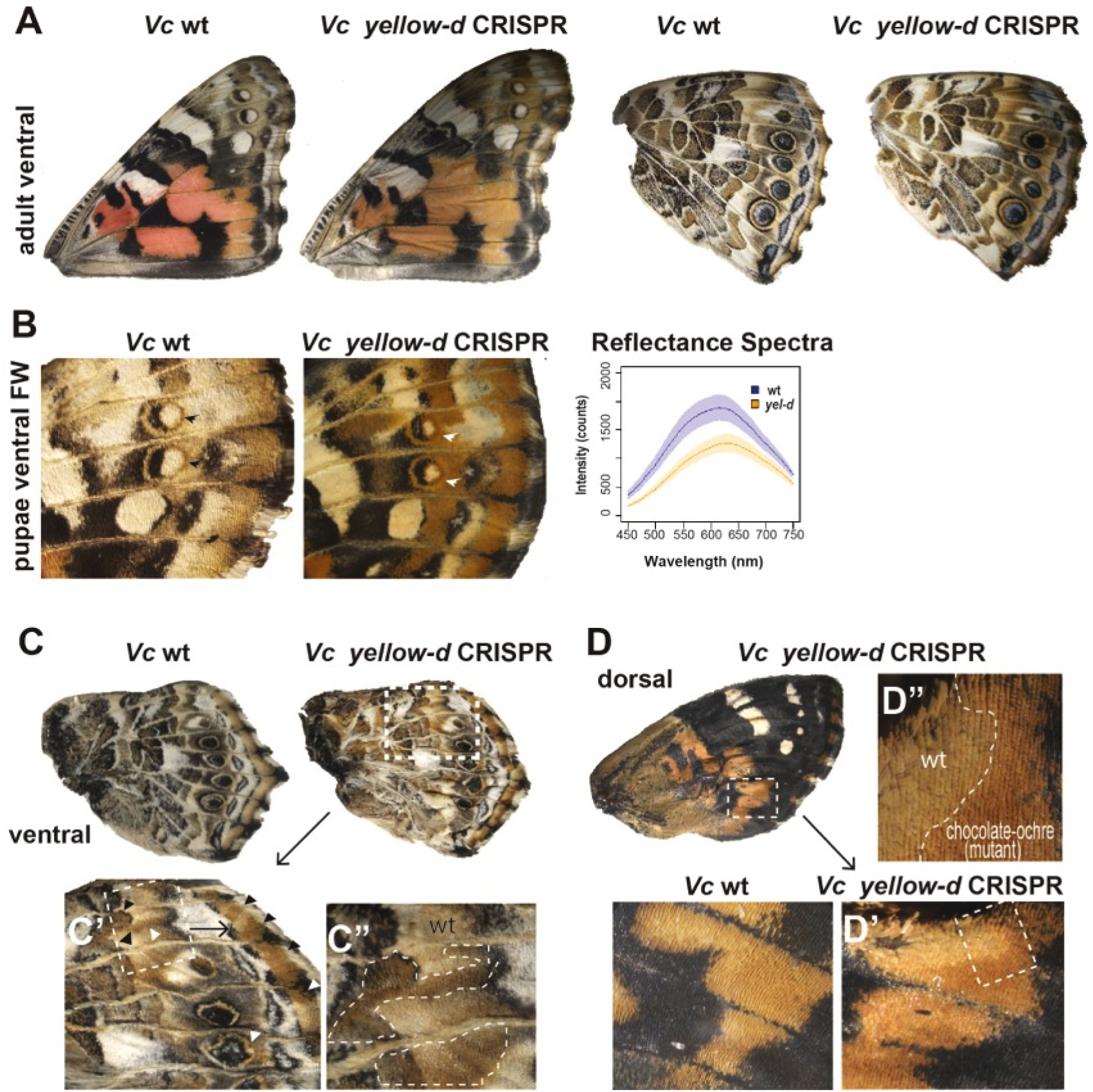
**Wing pattern-specific effects of *yellow-d* loss-of-function in *V. cardui*. (A)** Ventral view of wild type and *yellow-d* deletion wings in *V. cardui* shows promoted orange-brown pigmentation. (**B**) *yellow-d* deletion in *V. cardui* results in ectopic yellow pigments. Graphs of reflectance spectra of ventral forewing (white arrowhead) show significant shift in coloration between wild type and CRISPR mutants. (**C-C’’**) Ventral hindwing view of *yellow-d* deletion mosaic clones showing mutant ochre pigmentation in normally brown and grey color patterns. (**D-D’’**) *yellow-d* loss of function mosaic clones showing increased ochre pigmentation in dorsal forewing red patterns.

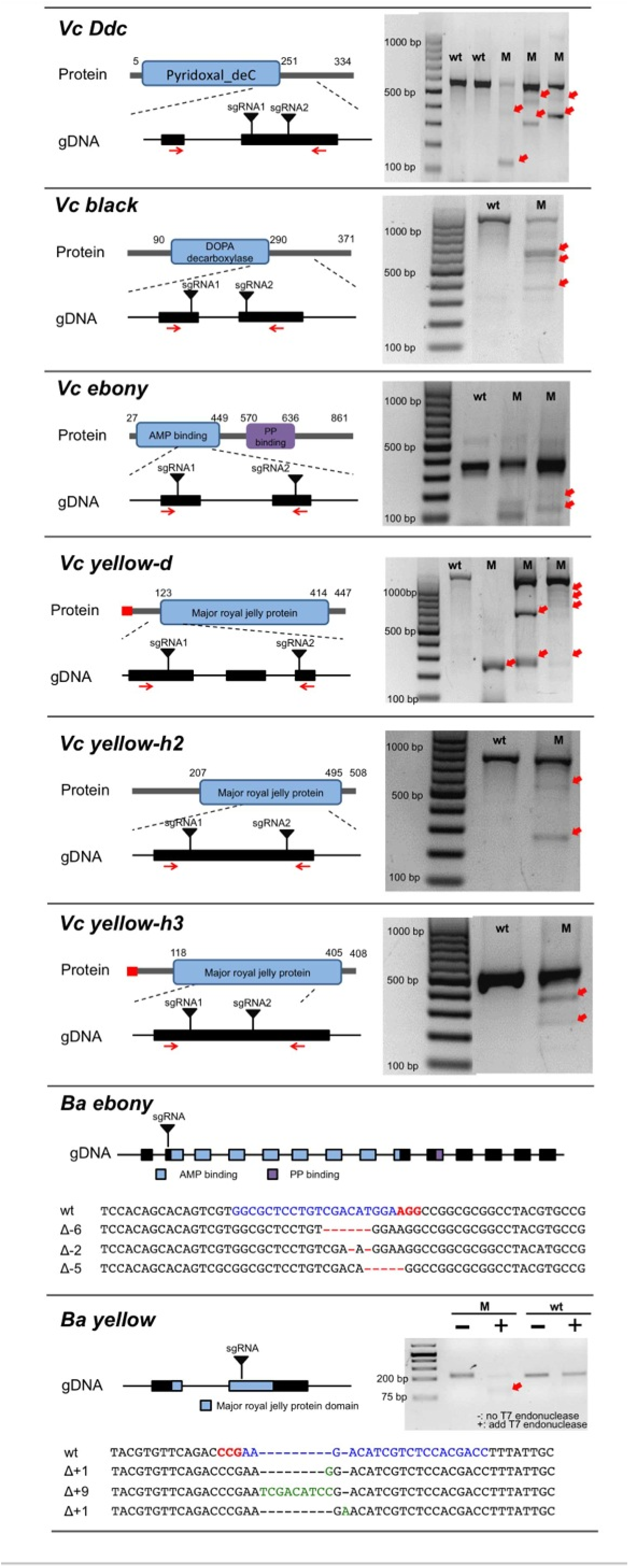

The RNA-seq and genome editing results in this study indicate previously undescribed genetic mechanisms underlying pigment development in butterflies. It is important to point out, however, that even though our approach is quick and powerful, use of G_0_ mosaics does have some limitations. One limitation that is general to all somatic mosaic studies, especially in smaller organisms such as insects, is that it is challenging to characterize specific causative alleles in specific clones. This is especially difficult in adult butterfly wings where wing scales are dead cuticle with degraded genomic DNA. Because of this our genotyping is limited to non-wing tissue, and thus primarily serves to confirm the accuracy and efficiency of our sgRNAs. Further, since we cannot perform molecular genotyping on specific wing clones we cannot confirm whether a clone is monoallelic or biallelic for induced lesions, and thus we can only speculate about potential dosage effects. Even despite these limitations, however, the easy-to-visualize nature of butterfly wing pigment phenotypes here allowed us to unambiguously validate the roles of multiple pigmentation genes in wing pattern development. Some gene functions we found to be deeply conserved in insects, while others we found to play novel or expanded roles in butterfly wing coloration. Understanding the genetic basis of adaptive features is one of the key goals in evolutionary and developmental biology, and our results provide a foundation for future work on the development and evolution of butterflies and other organisms as well.

## Author Contribution

R.D.R., L.Z., Ar.M, M.W.P., and Y.M. conceived the experiments. L.Z. performed RNA-seq work and data analysis. Genome editing and associated genotyping were performed by L.Z. (*V. cardui*), Ar.M and K.R.L.B. (*J. coenia*), Y.M. (*B. anynana*), M.W.P. (*V. cardui*, *P. xuthus*). R.D.R. and An.M. supervised experiments. L.Z., Ar.M., and R.D.R. wrote the manuscript with input from all co-authors.

## Acknowledgments

This work was supported by NSF grants IOS-1354318 and IOS-1557443 to R.D.R. and Ministry of Singapore grant MOE2015-T2-2-159 to An.M. We thank Joseph Fetcho and Joe DiPietro at Cornell University for assistance with microinjection, Kyle DeMarr and Benjamin John Brack for help raising butterflies, and Ellis Loew for help with microspectrophotometry. Finally, we are indebted to members of the Reed and Monteiro Labs as well as to Nipam Patel and his team for experimental assistance and discussions.

## References

Andersen, S. O., 2010 Insect cuticular sclerotization: a review. Insect Biochemistry And Molecular Biology 40: 166–178.

Arakane, Y., N. T. Dittmer, Y. Tomoyasu, K. J. Kramer, S. Muthukrishnan et al., 2010 Identification, mRNA expression and functional analysis of several yellow family genes in Tribolium castaneum. Insect biochemistry and molecular biology 40: 259–266.

Bassett, A., and J.-L. Liu, 2014 CRISPR/Cas9 mediated genome engineering in *Drosophila*. Methods 69: 128–136.

Bassett, A. R., C. Tibbit, C. P. Ponting and J.-L. Liu, 2013 Highly efficient targeted mutagenesis of *Drosophila* with the CRISPR/Cas9 system. Cell Reports 4: 220–228.

Beldade, P., and P. M. Brakefield, 2002 The genetics and evo–devo of butterfly wing patterns. Nature Reviews Genetics 3: 442–452.

Beldade, P., and C. M. Peralta, 2017 Developmental and evolutionary mechanisms shaping butterfly eyespots. Current Opinion in Insect Science 19: 22–29.

Biessmann, H., 1985 Molecular analysis of the yellow gene (y) region of *Drosophila melanogaster*. Proceedings of the National Academy of Sciences 82: 7369–7373.

Chen, S., P. Yang, F. Jiang, Y. Wei, Z. Ma et al., 2010 De novo analysis of transcriptome dynamics in the migratory locust during the development of phase traits. PLoS One 5: e15633.

Connahs, H., T. Rhen and R. B. Simmons, 2016 Transcriptome analysis of the painted lady butterfly, *Vanessa cardui* during wing color pattern development. BMC Genomics 17: 1.

Daniels, E. V., R. Murad, A. Mortazavi and R. D. Reed, 2014 Extensive transcriptional response associated with seasonal plasticity of butterfly wing patterns. Molecular Ecology 23: 6123–6134.

Ferguson, L. C., J. Green, A. Surridge and C. D. Jiggins, 2011a Evolution of the insect yellow genefamily. Molecular Biology And Evolution 28: 257–272.

Ferguson, L. C., L. Maroja and C. D. Jiggins, 2011b Convergent, modular expression of ebony and tan in the mimetic wing patterns of *Heliconius* butterflies. Development Genes and Evolution 221: 297–308.

Futahashi, R., and H. Fujiwara, 2005 Melanin-synthesis enzymes coregulate stage-specific larval cuticular markings in the swallowtail butterfly, Papilio xuthus. Development genes and evolution 215: 519–529.

Futahashi, R., J. Sato, Y. Meng, S. Okamoto, T. Daimon et al., 2008 yellow and ebony are the responsible genes for the larval color mutants of the silkworm Bombyx mori. Genetics 180: 1995–2005.

Futahashi, R., H. Shirataki, T. Narita, K. Mita and H. Fujiwara, 2012 Comprehensive microarray-based analysis for stage-specific larval camouflage pattern-associated genes in the swallowtail butterfly, *Papilio xuthus*. BMC Biology 10: 1.

Giraldo, M., and D. Stavenga, 2016 Brilliant iridescence of Morpho butterfly wing scales is due to both a thin film lower lamina and a multilayered upper lamina. Journal of Comparative Physiology A 202: 381–388.

Gorman, M. J., C. An and M. R. Kanost, 2007 Characterization of tyrosine hydroxylase from Manduca sexta. Insect biochemistry and molecular biology 37: 1327–1337.

Gorman, M. J., and Y. Arakane, 2010 Tyrosine hydroxylase is required for cuticle sclerotization and pigmentation in Tribolium castaneum. Insect biochemistry and molecular biology 40: 267–273.

Grabherr, M. G., B. J. Haas, M. Yassour, J. Z. Levin, D. A. Thompson et al., 2011 Full-length transcriptome assembly from RNA-Seq data without a reference genome. Nat Biotechnol 29: 644–652.

Guschin, D. Y., A. J. Waite, G. E. Katibah, J. C. Miller, M. C. Holmes et al., 2010 A rapid and general assay for monitoring endogenous gene modification. Engineered Zinc Finger Proteins: Methods and Protocols: 247–256.

Hines, H. M., R. Papa, M. Ruiz, A. Papanicolaou, C. Wang et al., 2012 Transcriptome analysis reveals novel patterning and pigmentation genes underlying *Heliconius* butterfly wing pattern variation. BMC Genomics 13: 288.

Hiruma, K., S. Matsumoto, A. Isogai and A. Suzuki, 1984 Control of ommochrome synthesis by both juvenile hormone and melanization hormone in the cabbage armyworm, *Mamestra brassicae*. Journal of Comparative Physiology B 154: 13–21.

Hori, M., K. Hiruma and L. M. Riddiford, 1984 Cuticular melanization in the tobacco hornworm larva. Insect Biochemistry 14: 267–274.

Ito, K., K. Kidokoro, S. Katsuma, T. Shimada, K. Yamamoto et al., 2012 Positional cloning of a gene responsible for the *cts* mutation of the silkworm, Bombyx mori. Genome 55: 493–504.

Koch, P. B., D. N. Keys, T. Rocheleau, K. Aronstein, M. Blackburn et al., 1998 Regulation of dopa decarboxylase expression during colour pattern formation in wild-type and melanic tiger swallowtail butterflies. Development 125: 2303–2313.

Kronforst, M. R., and R. Papa, 2015 The functional basis of wing patterning in Heliconius butterflies: the molecules behind mimicry. Genetics 200: 1–19.

Langmead, B., and S. L. Salzberg, 2012 Fast gapped-read alignment with Bowtie 2. Nature Methods 9: 357–359.

Lemieux, M. J., 2007 Eukaryotic major facilitator superfamily transporter modeling based on the prokaryotic GlpT crystal structure (Review). Molecular Membrane Biology 24: 333–341.

Li, X., D. Fan, W. Zhang, G. Liu, L. Zhang et al., 2015 Outbred genome sequencing and CRISPR/Cas9 gene editing in butterflies. Nature communications 6.

Liu, C., K. Yamamoto, T.-C. Cheng, K. Kadono-Okuda, J. Narukawa et al., 2010 Repression of tyrosine hydroxylase is responsible for the sex-linked chocolate mutation of the silkworm, Bombyx mori. Proceedings of the National Academy of Sciences 107: 12980–12985.

Liu, J., T. R. Lemonds, J. H. Marden and A. Popadić, 2016 A pathway analysis of melanin patterning in a Hemimetabolous insect. Genetics 203: 403–413.

Martin, A., K. J. McCulloch, N. H. Patel, A. D. Briscoe, L. E. Gilbert et al., 2014 Multiple recent co-options of Optix associated with novel traits in adaptive butterfly wing radiations. EvoDevo 5: 1.

Monteiro, A., 2015 Origin, development, and evolution of butterfly eyespots.

Morgan, T. H., 1916 Sex-linked inheritance in Drosophila. Carnegie institution of Washington.

Nijhout, H., and P. Koch, 1991 The distribution of radiolabeled pigment precursors in the wing patterns of nymphalid butterflies. The Journal of research on the Lepidoptera.

Nijhout, H. F., 1991 The development and evolution of butterfly wing patterns. Smithsonian series in comparative evolutionary biology (USA).

Osanai-Futahashi, M., K.-i. Tatematsu, K. Yamamoto, J. Narukawa, K. Uchino et al., 2012 Identification of the *Bombyx red egg* gene reveals involvement of a novel transporter family gene in late steps of the insect ommochrome biosynthesis pathway. Journal of Biological Chemistry 287: 17706–17714.

Pérez, M. M., J. Schachter, J. Berni and L. A. Quesada-Allué, 2010 The enzyme NBAD-synthase plays diverse roles during the life cycle of *Drosophila melanogaster*. Journal of Insect Physiology 56: 8–13.

Perry, M., M. Kinoshita, G. Saldi, L. Huo, K. Arikawa et al., 2016 Molecular logic behind the three-way stochastic choices that expand butterfly colour vision. Nature 535: 280–284.

Reed, R. D., and L. M. Nagy, 2005 Evolutionary redeployment of a biosynthetic module: expression of eye pigment genes *vermilion, cinnabar*, and *white* in butterfly wing development. Evolution & Development 7: 301–311.

Reed, R. D., R. Papa, A. Martin, H. M. Hines, B. A. Counterman et al., 2011 *optix* drives the repeated convergent evolution of butterfly wing pattern mimicry. Science 333: 1137–1141.

Robinson, M. D., D. J. McCarthy and G. K. Smyth, 2010 edgeR: a Bioconductor package for differential expression analysis of digital gene expression data. Bioinformatics 26: 139–140.

Roseland, C. R., K. J. Kramer and T. L. Hopkins, 1987 Cuticular strength and pigmentation of rust-red and black strains of *Tribolium castaneum*: Correlation with catecholamine and β-alanine content. Insect Biochemistry 17: 21–28.

True, J. R., K. A. Edwards, D. Yamamoto and S. B. Carroll, 1999 Drosophila wing melanin patterns form by vein-dependent elaboration of enzymatic prepatterns. Current Biology 9: 1382–1391.

Wallbank, R. W., S. W. Baxter, C. Pardo-Diaz, J. J. Hanly, S. H. Martin et al., 2016 Evolutionary novelty in a butterfly wing pattern through enhancer shuffling. PLoS Biol 14: e1002353.

Wittkopp, P. J., J. R. True and S. B. Carroll, 2002a Reciprocal functions of the *Drosophila* Yellow and Ebony proteins in the development and evolution of pigment patterns. Development 129: 1849–1858.

Wittkopp, P. J., K. Vaccaro and S. B. Carroll, 2002b Evolution of *yellow* gene regulation and pigmentation in Drosophila. Current Biology 12: 1547–1556.

Wright, T. R., 1987 The genetics of biogenic amine metabolism, sclerotization, and melanization in *Drosophila melanogaster*. Advances in Genetics 24: 127.

Wright, T. R., G. C. Bewley and A. F. Sherald, 1976a The genetics of dopa decarboxylase in *Drosophila melanogaster*. II. Isolation and characterization of dopa-decarboxylase-deficient mutants and their relationship to the α-methyl-dopa-hypersensitive mutants. Genetics 84: 287–310.

Wright, T. R., R. B. Hodgetts and A. F. Sherald, 1976b The genetics of dopa decarboxylase in *Drosophila melanogaster* I. Isolation and characterization of deficiencies that delete the dopa-decarboxylase-dosage-sensitive region and the α-methyl-dopa-hypersensitive locus. Genetics 84: 267–285.

Xia, A.-H., Q.-X. Zhou, L.-L. Yu, W.-G. Li, Y.-Z. Yi et al., 2006 Identification and analysis of YELLOW protein family genes in the silkworm, *Bombyx mori*. BMC Genomics 7: 195.

Zhang, L., and R. D. Reed, 2016 Genome editing in butterflies reveals that *spalt* promotes and *Distal-less* represses eyespot colour patterns. Nature communications 7.

Zhao, Y., H. Zhang, Z. Li, J. Duan, J. Jiang et al., 2012 A major facilitator superfamily protein participates in the reddish brown pigmentation in *Bombyx mori*. Journal of Insect Physiology 58: 1397–1405.

